# The Mechanical Power of Protein Folding

**DOI:** 10.1101/383711

**Authors:** Edward C. Eckels, Shubhasis Haldar, Rafael Tapia-Rojo, Jaime Andres Rivas Pardo, Julio M. Fernández

## Abstract

The delivery of mechanical power, a crucial component of animal motion, is constrained by the universal compromise between force and velocity of its constituent molecular systems. Here we demonstrate a switchable power amplifier in an Ig domain of the massive muscle protein titin. Titin is composed of many tandem repeats of individually foldable Ig domains, which unfold and extend during muscle stretch and readily refold when the force on titin is quenched during a contraction. Cryptic cysteine residues are common in elastic proteins like titin where they can oxidize to form intra-domain disulfide bonds, limiting the extensibility of an unfolding domain. However, the functional significance of disulfide-bonds in titin Ig domains remains unknown and may be fundamental to muscle mechanics. Here we use ultra-stable magnetic tweezers force spectroscopy to study the elasticity of a disulfide bonded modular titin protein operating in the physiological range, with the ability to control the oxidation state of the protein in real time using both organic reagents and oxidoreductase enzymes. We show that presence of an oxidized disulfide bond allows the parent Ig domain to fold at much higher forces, shifting the midpoint folding probability from 4.0 pN to 12.8 pN after formation. The presence of disulfide bonds in titin regulates the power output of protein folding in an all-or-none manner, providing for example at 6.0 pN, a boost from 0 to 6,000 zeptowatts upon oxidation. At this same force, single molecular motors such as myosin are typically stalled and perform little to no work. We further demonstrate that protein disulfide isomerase (PDI) readily reintroduces disulfide bonds into unfolded titin Ig domains, an important mechanism for titin which operates under a resting force of several pN *in vivo*. Our results demonstrate, for the first time, the functional significance of disulfide bonds as potent power amplifiers in titin and provide evidence that protein folding can generate substantial amounts of power to supplement the myosin motors during a contraction.

## Introduction

Mechanical systems are defined by their force-velocity relationship, which relates the magnitude of the work performed and the rate at which it is delivered. The inherent tradeoff between force and velocity holds true on scales ranging from man-made machines down to single molecule actuators and determines the peak power that can be delivered by a mechanical system (Ilton et al., 2018; Mahadevan and Matsudaira, 2000). In biological organisms, the magnitude of the mechanical power delivered determines the timescales on which it can respond to its environment, for example to evade a predator or to capture a prey. For these reasons, evolution has driven biological systems to optimize their power delivery. We know of many molecular scale actuators such as motors (Finer et al., 1994; Gelles et al., 1988; Howard, 1997), filament torsion (Sun et al., 1997; Way et al., 1995), filament assembly (Forscher et al., 1992; Theriot et al., 1992) and filament disassembly (Merz et al., 2000) that can deliver surprisingly large amounts of mechanical power on the scale of about ~1000 zW per molecular subunit. Mechanical power scales in networks of these molecules, reaching levels that are compatible with the environment of the organism (Mahadevan and Matsudaira, 2000). Thus, delivery of mechanical power is a central issue in understanding biological design, but very little is known about what dictates the mechanics at a molecular level in these power delivery systems.

One important addition was the discovery that protein folding does a large amount of mechanical work, on par with that measured in other molecular systems (Fernandez and Li, 2004; Rivas-Pardo et al., 2016). However, the force-velocity relationship for protein folding has never been measured and the power capabilities of a folding-driven molecular actuator remain unknown. Titin, the giant elastic protein of muscle tissues is a likely candidate for being an important source of mechanical power derived from folding (Eckels et al., 2018; Rivas-Pardo et al., 2016). While it is generally accepted that folding of a single protein domain can generate more work than the power stroke of a molecular motor, it remains unknown if protein folding can reach similar levels of power output as the motors in a muscle sarcomere (Bianco et al., 2016). For titin folding to contribute some energy during a muscle contraction requires that i) the magnitude of the work done by on the same order or greater than the myosin motors, ii) the work be delivered fast enough to provide a substantial part of the power output of a contracting sarcomere, iii) titin folding proceed fast enough that the myosin motors do not cause titin folding to go slack. While we have measured part (i) in our previous work (Rivas-Pardo et al., 2016), the final two questions can only be answered through determination of the force-velocity relationship for titin folding.

Here we use ultra-fast single molecule magnetic tweezers to measure, for the first time, the power output from titin protein folding. We demonstrate that oxidation of a single disulfide bond in the core of the protein amplifies the power output in an all-or-none manner, thus acting as a power switch. The disulfide bond makes this possible by raising the folding forces of the protein by more than eight piconewtons. This is likely an important mechanism in muscle for controlling the folding energetics of titin, which contains an abundance of cysteine residues shown to form disulfide bonds (Giganti et al., 2018; Manteca et al., 2017; Mayans et al., 2001). However, because titin operates under a constitutive resting force of several pN *in vivo*, enough to unfold and extend many Ig domains, it is unclear if there exists a mechanism for disulfide oxidation. We provide evidence that a typical oxidoreductase enzyme, protein disulfide isomerase (PDI), is able to reversibly induce disulfide formation at forces as high as 5 pN, and possesses additional chaperone activity to assist folding. These data provide concrete evidence that the speed and power output of titin protein folding could indeed contribute to the energetics of a muscle contraction. While much weight has been given to equivalent force spectroscopy experiments on the muscle myosin II motors, in understanding the scaling of force generation to tissue levels, the current models will need to re-evaluate the relative power contributions of titin and the myosin motor given the data presented here.

## Results

### Real time control of single Ig domain oxidation status

The magnetic tweezers instrument provides a natural way for probing the force-velocity relationship of a nanomechanical system because it supplies feedback-free force-clamp along with high resolution tracking of folding trajectories. Here we measure the force-velocity relationship of a tandem modular protein construct containing eight repeats of cardiac titin Ig domain I27 with two cysteines that can form a single disulfide bond (Wiita et al., 2006). The eight identical repeats of the I27 domain in the expressed construct generate an unambiguous fingerprint upon unfolding. In the oxidized form (I27^OXD^), the formed disulfide limits the extension of the amino acid backbone upon unfolding (Figure 1A), resulting in 11 nm upwards steps as each Ig domain unfolds up to its disulfide bond (Figure 1B, blue trace) during the *extend* pulse. In contrast, the same Ig domains with the reduced disulfide (I27^RED^) demonstrate 25 nm upwards steps, identical to those observed in the wild-type I27 domain (Figure 1B, red trace). After unfolding, the force is reduced to allow the polypeptide backbone to collapse during the *refold* pulse. There is a marked difference in the folding trajectories of the reduced and oxidized Ig domains – the reduced Ig domains demonstrate a gradual downward stepwise trajectory, a combination of six downward events and two upwards events, indicating a net total of four folding events. By pulling again at high force in the *probe* pulse, the four 25 nm unfolding steps confirm the refolding count. By contrast, the oxidized Ig domain shows immediate collapse during the refolding pulse and eight 11 nm steps during the probe pulse, demonstrating complete refolding.

**Figure 1:**
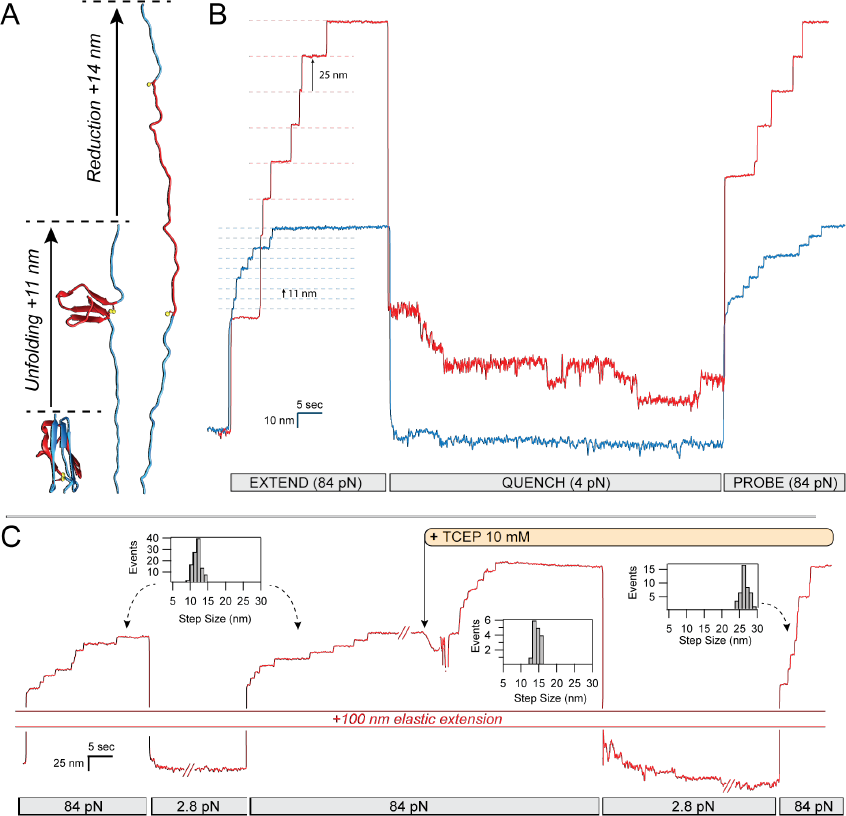
Real-time control of cryptic disulfide bonds in single polyproteins under force. **(A)** Under force, an Ig domain extends up to its buried disulfide bond, exposing it. Upon reduction, a further extension occurs. **(B)** The oxidation status and effect of cryptic disulfide bonds are evident in these two traces of polyproteins containing eight repeats of a titin Ig domain. Oxidized polyproteins extending at a high force of 84 pN increase their length in steps of 11 nm (blue trace), whereas reduced polyproteins extend in steps of 25 nm (red trace). While the rate of folding of the reduced polyprotein at 4 pN is slow, oxidation greatly accelerates the folding rate, driving it to completion at the same force. The increased folding probability at 4 pN is measured from the number of steps recovered with a probe pulse at 84 pN (red trace 4, blue trace 8). **(C)** The status of a single polyprotein can be easily changed from oxidized to reduced by introducing a reducing agent into the solution (TCEP), while the polyprotein is fully unfolded, exposing its thiols to the solution. Step size histograms record the changes observed from extending a fully oxidized polyprotein (11.4 +/‐ 2.1 nm); immediately after addition of TCEP (14.6 +/‐ 1.6 nm) and after refolding the resulting fully reduced polyprotein (25.6 +/‐ 2.2 nm).

The oxidation status of the disulfide bond can be controlled in real-time as depicted in the chart-recorder style single molecule trace in Figure 1C. In this experiment, the molecule is first stretched at high force (84 pN) to unfold eight oxidized Ig domains, each with a step size of 11 nm (inset histogram, Fig. 1C). The force is quenched to 2.8 pN to allow for refolding while a buffer containing 10 mM tris(2-carboxyethyl) phosphine (TCEP) is prepared for exchange into the chamber. Unfolding the molecule again at high force demonstrates eight upwards steps of 11 nm, after which the reducing buffer is introduced through three successive washes of 100 uL, a volume ten-fold higher than the volume contained inside the flow cell. Reduction by TCEP leads to eight successive 14 nm steps caused by cleavage of the disulfide bond and release of the cryptic length sequestered behind the bond (Figure 1A). With the reducing agent still present in the solution, the force on the molecule is again quenched to 2.8 pN to allow for refolding. A final probe pulse at 84 pN demonstrates eight 25 nm steps – indicating complete refolding of the protein in its reduced state.

### A single disulfide bond shifts titin folding to higher forces

This assay was used to determine the folding probability of the oxidized versus reduced Ig domain over a wide range of forces, calculated as described previously (Rivas-Pardo et al., 2016). After unfolding the I27^OXD^ construct using an *extend* pulse, the refolding force was clamped at a force ranging from 5-18 pN for 100 s, after which the *probe* pulse revealed how many domains had folded (Figure 2A). The I27^OXD^ domains experienced very little refolding at 15.3 pN, occasionally one domain per octamer, but exhibit 7 folding events at a refolding force of 10.6 pN. The step sizes observed during the refolding pulse scale according to the freely-jointed chain (FJC) polymer model (Flory, 1953) and match the previously reported value for contour length (Ainavarapu et al., 2007), 12.7 nm, with a slightly larger Kuhn length of 0.65 nm that is a better fit in the low force regime explored by the magnetic tweezers (Figure 2C). After obtaining the data from the oxidized domain, the protein was reduced with 10 mM TCEP, which was kept in solution to prevent spontaneous re-oxidation or unwanted side reactions of the two reduced cysteine residues while measuring the folding probability. The I27^RED^ construct showed dramatically different folding trajectories, requiring forces as low as 5 pN to begin refolding, and forces down to 2.5 pN for complete refolding of all eight domains (Figure 2B). These size of refolding events also demonstrated scaling with the FJC model, with a Kuhn length of 1.15 nm, and a contour length of 28.4 nm (Figure 1C, red curve). The folding probability as a function of force demonstrates a large, 8.4 pN shift in the midpoint of the folding probability according to the fit with a sigmoid curve (Supplementary Methods). This shift is clearly demonstrated by the raw data, which shows rapid refolding of the I27^OXD^ domains at a force of 10.6 pN, but rather slow refolding from I27^RED^ at a force of 4.3 pN (Figure 2A vs 2B). The cause of this increase in the folding kinetics and lowering of the refolding barrier may arise from either the reduced entropy of the backbone imposed by the presence of the disulfide bond or perhaps some residual structure in the peptide loop delimited by the disulfide bond that serves to nucleate folding.

**Figure 2:**
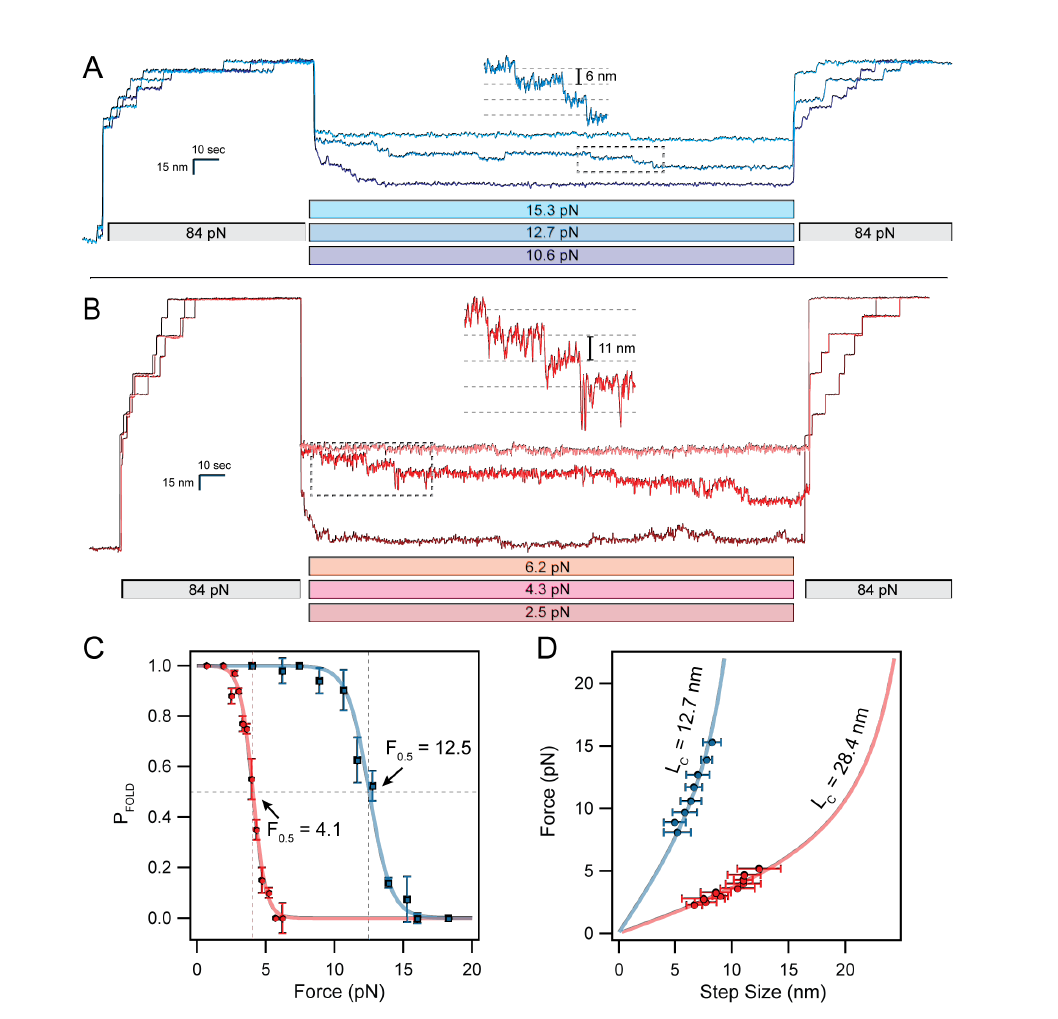
Cryptic disulfide bonds alter the folding dynamics of titin Ig domains. **(A)** After full extension, oxidized polyproteins are allowed to fold by quenching the force to 10-16 pN for 100 sec. While only a single folding event is observed at 16 pN, complete refolding occurs at 10.6 pN. **(B)** Reduced polyproteins fold at a much lower range of forces in the 2-5 pN range. **(C)** The folding probability is calculated as the number of refolded domains (probe pulse) divided by eight. Oxidation shifts the midpoint folding probability from 4.1 pN to 12.5 pN. The data was collected from several different molecules, with n=11 for I27^OXD^ and n=9 for I27^RED^ **(D)** The folding steps observed during the quench pulse (insets in A and B) scale in size with the applied force. Oxidized folding steps (blue circles) are fit by the FJC model (solid blue line, see supplementary equations) using a contour length of 12.7 nm. Reduced folding steps (red circles) are fit using a much larger contour length of 28.4 nm (solid red line). These data came from the same molecules measured in panel (C).

### Protein Disulfide Isomerase (PDI) mediates disulfide reformation under force

In vivo, disulfide bonds are most commonly introduced via a specialized set of electron shuttling enzymes called oxidoreductases. The most common of these enzymes in mammals is protein disulfide isomerase (PDI) which consists of four domains with thioredoxin-type folds. Two of the domains, A‐ and B-, contain a catalytic *CXXC* motif that can cleave and reform disulfide bonds in substrate proteins, as well as two other domains (A’ and B’) lacking the catalytic motif, thought to possess chaperone activity (Gruber et al., 2006). It is not clear whether or not the catalytic domains also possess chaperone activity. The magnetic tweezers was recently used to demonstrate that Trigger Factor, a prokaryotic chaperone, is capable of accelerating folding in substrates under force, which may be a common scenario with polypeptides emerging from the ribosome, the translocon pore, and other molecular tunnels (Haldar et al., 2017). Titin is another case of a protein that is naturally stretched *in vivo*, and experiences a constitutive resting force due to the geometry of the sarcomere (Linke, 2018; Llewellyn et al., 2008). It is unclear if the oxidoreductase machinery of the cell is capable of functioning to correctly oxidize disulfide bonds in substrates that experience such forces, but there is evidence that PDI plays an important protective role in cardiac muscle tissue (Severino et al., 2007). Here we examine the effects of the single catalytic PDI A-domain (referred to as PDI) on the re-oxidation and refolding of the I27 substrate.

After unfolding all eight I27^OXD^ domains, 60 uM PDI^RED^ was flowed into the chamber to react with the exposed substrate disulfide bonds (Figure 3A). Cleavage of substrate disulfide was marked by eight 14.1 nm upwards steps, identical to those seen in the presence of TCEP, with the exception that >200-fold less concentration was used, indicating the highly nucleophilic nature of the catalytic PDI thiols. Upon complete cleavage of the disulfide bonds, the force was reduced to 5 pN to allow the protein to fold. Unlike the situation with TCEP where little to no folding occurs at 5 pN, there is a downwards staircase of seven steps, indicating refolding. A subsequent probe pulse of 84 pN reveals both 11‐ and 14‐ nm steps, indicating that both disulfide reformation and refolding have occurred. In the cellular environment, PDI is maintained in its oxidized state with the help of several other electron shuttling enzymes. In order to recreate such an environment, the same assay was repeated with oxidized PDI (see Methods) and reduced I27 (Figure 3B). I27^OXD^ was first unfolded (Fig 3B, part 1) and reduced through addition of 10 mM TCEP (Fig 3B, part 2), after which the force was reduced to 5 pN. This confirms that there is little folding activity at 5 pN, as indicated by the single reduced (25 nm) step observed on the following probe pulse. Next the TCEP was thoroughly removed from the chamber with five washes of 100 uL buffer. Reducing the force to 2 pN and pulling again shows the presence of seven I27^RED^ domains (Fig 3B, part 3). Finally, 60 uM PDI^OXD^ was added to the flow chamber through two washes to create an oxidizing environment. Reduction of the force to 5 pN again shows seven downwards steps, in stark contrast with the folding trajectory in the presence of TCEP. The last probe pulse demonstrates seven each of 11‐ and 14‐ nm steps, indicating re-oxidation of the disulfide bond in addition to refolding (Fig 3B, part 4)

**Figure 3:**
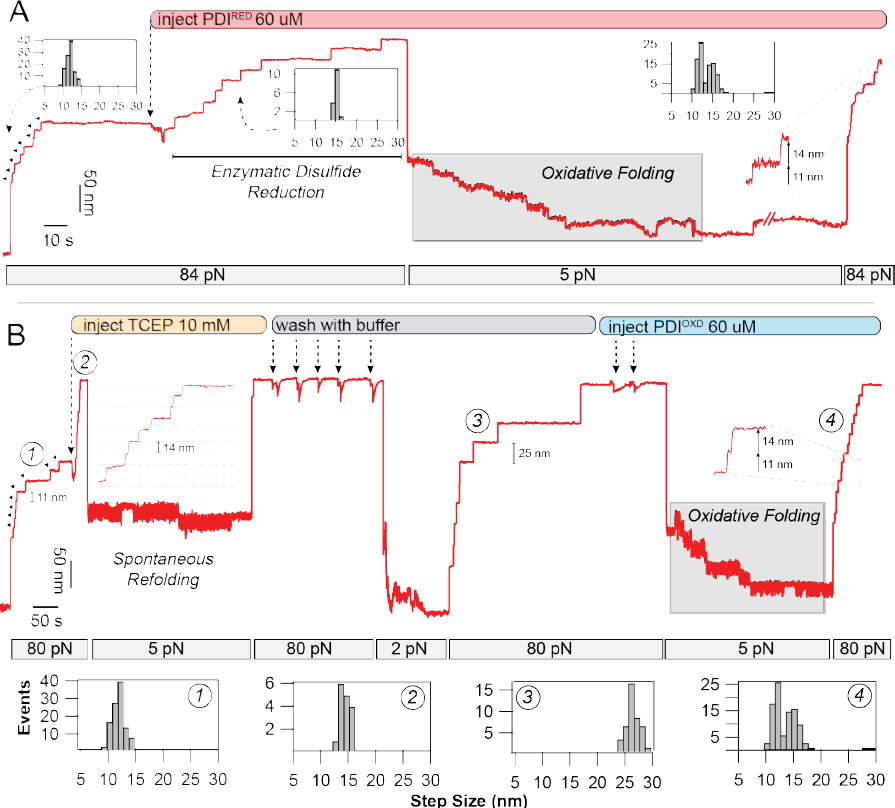
Oxidoreductase mediated disulfide cleavage and reformation under force. **(A)** After full unfolding of eight I27ox domains (11.4 nm steps), PDI^RED^ was added to the flow cell, causing eight distinct disulfide cleavage events (14.6 nm steps). After quenching the force to 5 pN, a downward folding staircase is observed. Subsequent unfolding at 84 pN reveals a mixture of 11‐ and 14‐ nm steps, indicating reformation of the disulfide bond. **(B)** Disulfide reformation could also be achieved with oxidized PDI: after full unfolding of eight I27^OXD^ domains, TCEP (10 mM) was added to the flow cell, resulting in eight 14.6 nm steps due to reduction of the disulfide bond. Quenching the force to 5 pN resulted in only a single refolding event. The protein was stretched again at high force to remove the TCEP with 5x washes using buffer. Reducing the force to 2 pN caused refolding of 7x I27^RED^ domains, as indicated by the 25 nm steps. After full unfolding, PDI^OXD^ was added to the flow cell, and the force was reduced to 5 pN again. In the presence of the oxidoreductase, a folding staircase is again observed. Subsequent pulling at 84 pN demonstrates 7x 11‐ and 14‐ nm steps, indicating nearly complete reformation of the disulfide bond at 5 pN.

### The catalytic domain of PDI possesses chaperone activity

The assay, utilizing I27^OXD^ and PDI^RED^, allowed for many cycles of enzymatic reduction and oxidative folding to be conducted at different refolding forces (Figure 4A). Scanning the refolding forces from 3.5 to 6.5 pN permitted direct observation of the folding trajectories in the presence of PDI and the ability to control the amount of refolding. The folding probability, calculated as the number of 11‐ and 25‐ nm steps in the probe pulse, divided by the number of 11‐ and 25‐ nm steps on the extend pulse, shows a clear shift of the sigmoidal folding probability to higher forces (5.0 pN midpoint) in the presence of PDI (Figure 4B). We attribute this to the mixed disulfide formed between PDI and I27 after cleavage of the I27^OXD^ internal disulfide bond, which increases the propensity of the substrate to fold under force. By contrast, folding in the presence of TCEP, which does not form stable adducts with protein thiols, has a midpoint folding force of 4.0 pN. The step-sizes measured at each force were collected and averaged and found to fall directly on the FJC curve of folding step sizes when the I27 was maintained in the reduced state by TCEP. For each folding step observed during the quench pulse, there is a corresponding re-oxidized (11 nm) unfolding event. Failure to reform the disulfide bond after folding was exceedingly rare. Taken together, these data suggest that PDI re-introduces the disulfide bond after the first step of protein folding, collapse of the polypeptide backbone to a high entropy molten globule state. This confirms the conclusions, that protein folding drives disulfide formation, inferred from previous single molecule data that lacked resolution to resolve the individual folding steps (Kahn et al., 2015; Kosuri et al., 2012). Furthermore the shift in folding probability to higher forces suggests a *bona fide* chaperone activity of the PDI domain with extended polypeptides, indicating a pathway for disulfide bond introduction into titin even under the resting forces it experiences in muscle.

**Figure 4:**
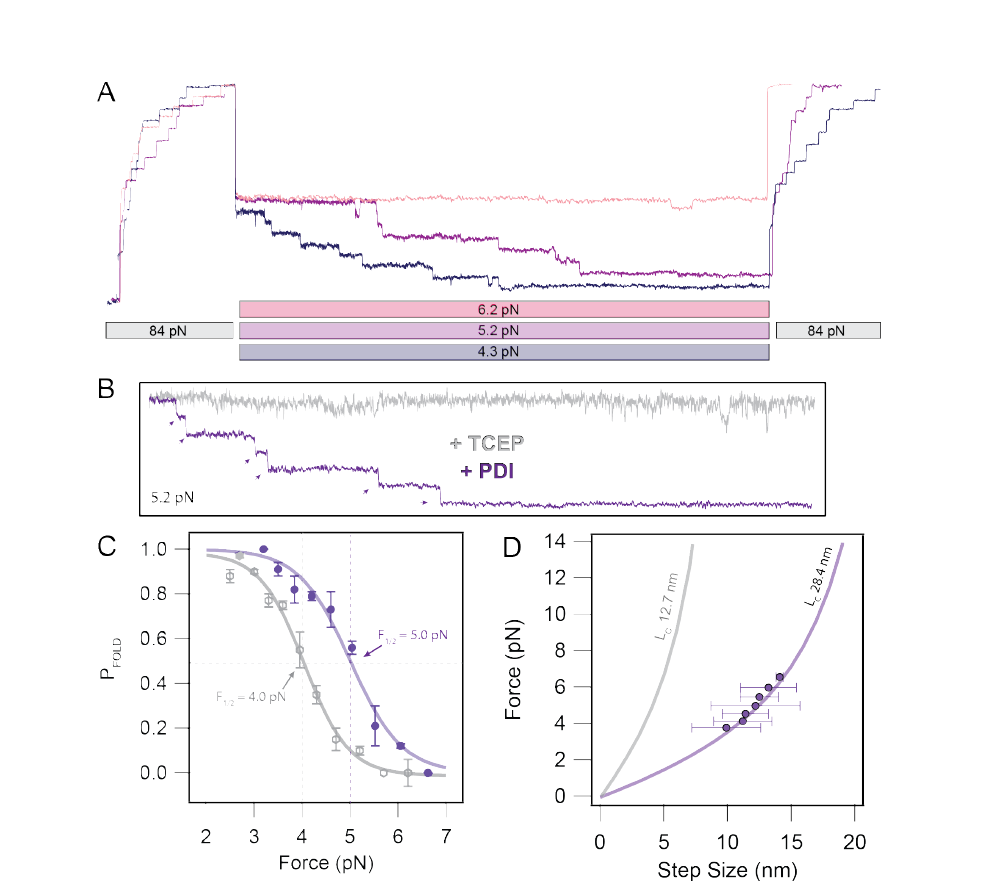
PDI introduces disulfide bonds after accelerated polypeptide collapse. **(A)** Folding trajectories of I27 in the presence of 60 uM PDI^RED^ at 6.2, 5.2, and 4.3 pN. Both the extend and probe pulses contain both 11‐ and 14‐ nm steps due to initial cleavage of the disulfide bond and subsequent oxidoreductase mediated reformation of the disulfide bond during the quench. **(B)** Comparison of individual folding trajectories of I27^RED^ in the presence of 10 mM TCEP versus I27^OX^ after cleavage by 60 uM PDI^RED^. The greatly favored folded state in the presence of PDI suggests chaperone activity from the oxidoreductase. **(C)** The folding probability of I27 is shifted in favor of the folded state in the presence of PDI. The sigmoidal fits demonstrate a 1.0 pN shift of the midpoint folding probability from 4.0 pN in the presence of TCEP to 5.0 pN in the presence of PDI^RED^. **(D)** Folding step sizes observed in the presence of PDI closely followed the FJC model for the I27^RED^ domain, even if there is disulfide reformation. This suggests that the oxidoreductase reintroduces the disulfide bond only after there is collapse of the polypeptide backbone.

### Disulfide formation boosts the work done by titin folding

Recent experimental data on muscle fibers suggests that the energetics of titin folding might be important for active muscle contraction in addition to passive mechanics. From the measurements presented so far, it is not immediately clear if presence of an extension-limiting disulfide bond will increase or decrease the mechanical work delivered by a folding Ig domain because higher folding forces and smaller step sizes have opposite effects on work delivery. The maximum work can be calculated simply as the product of the applied force and the mean step size at that force. However in the force range explored for titin folding, there is an equilibrium between folding and unfolding steps, so the net amount of work delivered by folding a single titin Ig domain is equal to the maximum work (empty squares, Figure 5) times the folding probability (empty diamonds, Figure 5). This quantity is called < *W* >, the expected value of the work output (Rivas-Pardo et al., 2016) (solid circles, Figure 3). The I27^RED^ folding process generates a peak of only 25.6 zJ at 4.2 pN whereas I27^OXD^ folding work peaks at a value of 63.9 zJ and 11.7 pN (Figure 5, solid curve). Oxidation of the single disulfide bond in this protein therefore doubles the magnitude of the expected work performed Ig domain folding. Although these work values are much higher than the average energy (~36 zJ) delivered by muscle motor proteins (Piazzesi et al., 2007), a better assessment of the relevance of protein folding on a physiological scale is the rate at which the energy is delivered.

**Figure 5:**
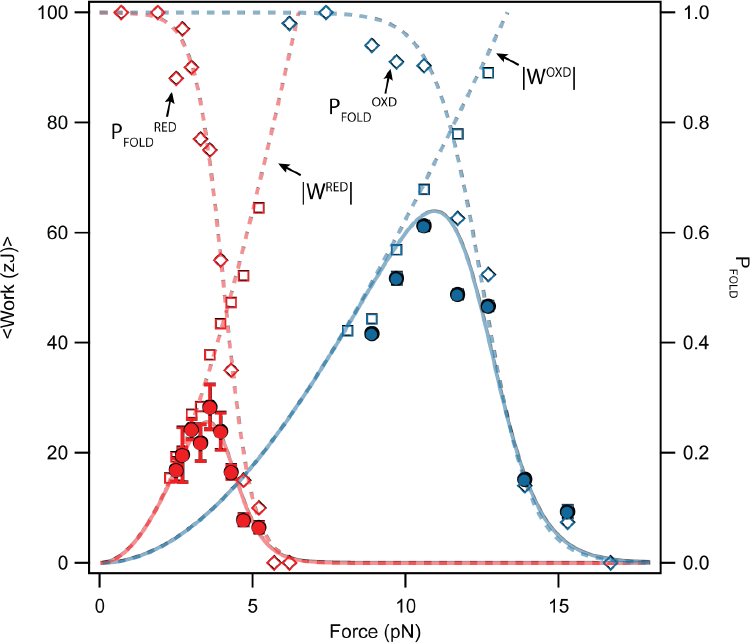
Oxidation increases the work done by protein folding and shifts its peak to higher forces. We calculate the expected value of the work delivered by folding steps as the product of folding work times the folding probability. The expected value of the folding work delivered by a reduced Ig domain peaks at force of 3.5 pN with a value of 25.6 zJ (solid red line), calculated by multiplying the reduced folding probability (empty red squares, dashed line) times the reduced folding work (empty red diamonds, dashed line). Oxidation causes large effects on the mechanical work delivered by protein folding: the expected value of the folding work delivered by an oxidized Ig domain is now 2.5 times bigger (63.9 zJ) and peaks at 11 pN (solid blue line; from the oxidized folding probability (empty blue diamonds, dashed line) times the oxidized folding work (empty blue squares, dashed line). However, in both cases, the delivery of the folding work at their peak is very slow due to measured folding kinetics.

### A model for power delivery from protein folding

During a muscle contraction, force development occurs within milliseconds after activation of the muscle fiber (Huxley and Simmons, 1971), redistributing some of the load from titin to the crossbridges themselves. Here we aim to mimic these force changes in titin using ultra-fast force quenches to determine if there is significant power output from titin on the timescales relevant in muscle (Figure 6A). After a high force *extend* pulse at 80 pN, the force was ramped down to 16 pN over the course of 0.5 seconds, where I27^OXD^ does not fold (Figure 2C). After settling at 16 pN for 1 second, the magnets were stepped as quickly as possible (~5 ms) to provide a force in the range of 2-10 pN. Tracking of the protein trajectory after these very fast force quenches permits a measurement of the power output due to protein folding. As seen in Figure 6A, the folding trajectories are so rapid that it is difficult to distinguish individual folding steps. The change in length at 16 pN before and after the force quench, as well as the presence of eight steps in the final probe pulse (not shown), indicate complete refolding of the protein construct. The force ranges over which the folding kinetics could be tracked were 3-10 pN for I27^OXD^ or 2-3 pN for I27^RED^. The lag introduced by changing the magnet position (< 5ms) and the amplitude of the folding contraction were the limiting factors at lower forces. The effective shortening rate during the refolding pulse was determined by fitting a single exponential to the trajectory (inset, Figure 4A). The amplitude of the exponential was fixed to eight times the step size as predicted by the freely-jointed chain model (Figure 2D), thereby excluding the entropic collapse of the polymer due to the change in force. The observed rate constants are plotted for both the oxidized and reduced forms of the protein are shown in Figure 4B, blue and red data points, respectively. The folding rates are fit with a modified exponential model that takes into account the free energy of a polypeptide (Chen et al., 2015; Dudko et al., 2008) modeled as a freely-jointed chain polymer extended up to the pulling force:

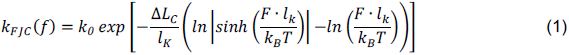

**Figure 6:**
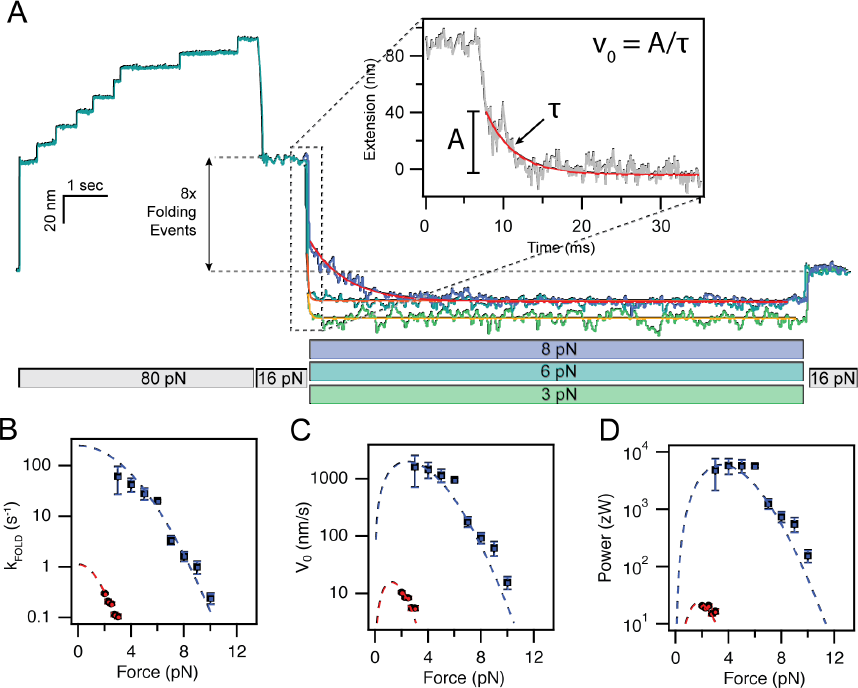
Oxidation acts as a power switch controlling the delivery of protein folding work. **(A)** Force protocol for measuring fast folding contractions at forces where the folding probability is always one. A fully extended oxidized polyprotein is first quenched to 16 pN, where no folding occurs. A rapid quench follows, resulting into a brief elastic recoil and a very fast folding contraction. A subsequent increase of the force up to 16 pN reveals a much shorter extension, marking the folding of all eight domains. The speed of the folding contraction increases greatly as the force of the quench is reduced. The folding contractions are fit with a single exponential to measure the folding rate. **(B)** Shows the force dependency of the folding rate for oxidized domains (blue squares) and reduced domains (red squares). The dashed lines correspond to fits using equation 1. **(C)** Force dependency of the initial velocity of the folding contraction (*V_0_*), measured as the amplitude of the contraction divided by its time constant (inset, panel A). The dashed lines correspond to fits from fits from equation 2. **(D)** the power delivered by the folding contraction is then calculated as *V_0_* · *F*. The dashed lines are fits of equation 3. The figure shows that oxidation of the polyprotein increases the peak output power of its folding contraction by more than 300 fold, reaching ~ 6000 zW. Data was averaged from n=5 molecules for I27^OXD^ and from n=3 molecules for I27^RED^.

Here Δ*L*_*C*_ is the Kuhn length as determined from the previous fits to the freely jointed chain (Figure 2D). The curvature of the folding rates due to movement of the folding transition state with force is well accounted for by this model (dashed fits, Figure 6B). These experiments are nearly identical to those performed on muscle fibers to determine the force-velocity relationship of muscle, where shortening velocities are recorded after a small step change in the load. The muscle fiber typically exhibits <100 nm shortening per half sarcomere over tens of milliseconds (Caremani et al., 2016; Linari et al., 2015; Piazzesi et al., 2007). Therefore we consider the initial shortening velocity, *ν*_0_, of a titin molecule as the physiologically relevant measurable here to mimic the equivalent muscle experiments. The velocity, *ν*_0_ slope of the folding trajectory at time zero, and is calculated from the amplitude of the exponential fit multiplied by the force dependent folding rate, given by

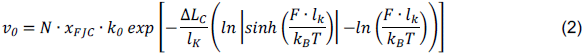

where *N* is the number of folding domains and *x*_FJC_ is the force dependent step size of folding calculated from the freely jointed chain. Unlike the force-velocity relationship of muscle, which does not have a maximum according to the hyperbolic Hill equation, the force-velocity relationship shown here is peaked due to the rapidly diminishing folding step size as the force approaches zero. Despite the contour length of the I27^OXD^ being less than half that of the reduced domain, the incredibly accelerated folding rate causes the folding velocity of I27^OXD^ to be ~300-fold higher at a force of 3 pN. From these data, the power output can be easily calculated by multiplying the applied force by the measured shortening velocity.

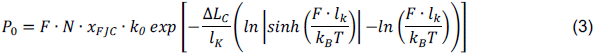

The power output of this disulfide bonded titin construct reaches much higher than any other protein ever observed, up to 5,870 zW measured at a pulling force of 4 pN, with a predicted peak power of 6,530 zW at a force of 3.6 pN according to the fit by equation (3). By contrast, the largest power output measured from the reduced domains reaches only to 21 zW at a force of 2.5 pN. This is comparable to, but slightly under the power output estimated from the wild type I27 and I10 domains (neither contain disulfide bonds), which reach approximately 200 zW. It is important to keep in mind that this protein has several different residues from the I27 used in previous studies, which is known to alter folding rates and folding forces. From these measurements it is clear that the disulfide bond acts as a switch that doubles the amount of work done by folding, but is able to boost the power output of folding by several orders of magnitude.

## Discussion

Cysteine residues are widely recognized as molecular switches because of their reversibly reactive nature (Littler et al., 2010; Müller et al., 2015), yet the role of disulfide bonds on the dynamics of mechanically active proteins in the physiological force range has never been explored due to instrumental limitations. Magnetic tweezers provide the ability to accurately control the applied forces in the piconewton range, with the additional advantage of being able to exchange buffers throughout the course of an experiment to alter the redox status of protein thiols. Here we use the magnetic tweezers to demonstrate that disulfide bonding is a switch-like regulator of the power output from Ig domain folding, converting a soft and extensible polypeptide into a potent power-delivering actuator. At a force of 6 pN, the reduced protein domain is completely unable to fold (Figure 2B and 7). Simple oxidation of the disulfide bond allows the protein to fold at the same force of 6 pN with a predicted step size of 4 nm, each step thus delivering 24 zJ of work. With an initial folding velocity of 1,000 nm/s, this equates to a power delivery of 6,000 zW (Figure 7). The switch like behavior reported here is easily tuned by the redox status of the muscle, which is determined by antioxidant and oxidoreductase levels, which change with exercise, stress, ischemia, and various homeostatic responses. We further demonstrate that a canonical oxidoreductase enzyme, protein disulfide isomerase, is able to introduce disulfide bonds into titin even when it is stretched by forces in the range thought the be experienced in a relaxed muscle. These experiments unexpectedly revealed a chaperone-like quality of PDI, shifting the folding to 1.0 pN higher forces.

**Figure 7.**
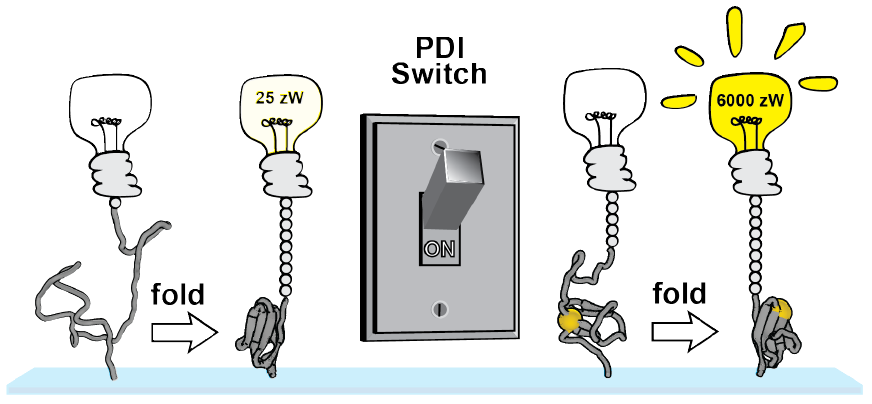
A PDI-mediated disulfide switch of the power output from titin folding. Upon initiation of a muscle contraction, the force on titin is lowered through engagement of the crossbridges. Titin domains lacking cysteines or containing reduced disulfide bonds are unable to fold rapidly and there for contribute very little peak power, reaching a maximum of only 25 zW at a force of 2 pN (left panel). Oxidation of titin disulfide bonds through the PDI machinery of the cell allows titin domains to fold at higher forces and more rapidly. Despite the collapse step being much larger in the case of the reduced domain, the forces and kinetics from folding of the oxidized domain at 4.0 pN are so much higher that the power output is two orders of magnitude greater.

Muscle tissue routinely experiences wide fluctuations in the redox balance of the cytosol especially during periods of high metabolic demand, for example exercise. It is highly likely that the abundant cysteine residues in titin and other sarcomeric proteins (myosin, actin, troponin C, ryanodine receptor, protein kinase C, myosin binding protein C) regulate muscle mechanics through cysteine chemistry (Steinberg, 2013). The nearly crystalline arrangement of filaments in the sarcomere allows for such dense protein packing that protein thiol concentrations can reach the millimolar range. It is therefore impossible for intracellular antioxidants to completely buffer protein thiols from redox reactions. The role of the abundant cysteines in titin has long remained unclear, and although it had been hypothesized that disulfide bonding (Kosuri et al., 2012; Wiita et al., 2007), disulfide isomerization (Alegre-Cebollada et al., 2011; Giganti et al., 2018), and other post-translational modifications (Alegre-Cebollada et al., 2014) might alter titin elasticity, never before had their role in power output been explored. Acquisition of disulfide bonds in the intracellular compartment can be achieved through several different mechanisms, including formation of intermediates with reactive oxygen species or low molecular weight disulfide compounds. While there is evidence that low molecular weight thiols can induce disulfide reformation in peptides at zero force (Beedle et al., 2016, 2017, 2018), this is unlikely a viable mechanism at the 4-10 pN stretching forces experienced by titin *in vivo*. A more likely mechanism for introduction of disulfide bonds is through oxidoreductase enzymes, which have evolved to recognize unfolded substrates containing cysteine residues. Indeed PDI is one of the genes upregulated in cardiomyocytes in response to myocardial infarction, and plays an important role in post-MI remodeling (Severino et al., 2007). Previous studies using PDI and I27 utilized AFM based force spectroscopy to measure the rates of cleavage and reformation of this same disulfide bond (Kosuri et al., 2012). However, the large fluctuations of the AFM cantilever at low forces prevented direct observation of the folding events. The magnetic tweezers overcomes this because by operating under highly damped, passive force clamp conditions, enabling accurate tuning of the force to fractions of a piconewton.

Refolding trajectories of I27 in the presence of PDI reaffirm the mechanism of PDI proposed by the previous AFM studies, namely that re-oxidation of the disulfide bond is preceded by entropic collapse of the I27 polypeptide into a molten globule like structure, as indicated by the step sizes of folding measured in Figure 4D. What could not be probed in the previous studies was the ability of PDI to bias the folding reaction to the native folded state. While it is intuitive that presence of a pre-formed disulfide bond allows I27 to fold at higher forces (12.5 pN vs. 4.0 pN midpoint), it is not immediately clear why having a mixed disulfide between PDI and I27 also shifts the folding probability of I27 to higher forces (Figure 3B and 4C). One possible explanation is the chaperone activity that is a supposedly generic property of the thioredoxin fold, independent of its catalytic activity (Quan et al., 1995). The catalytic *CXXC* motif is surrounded by a small hydrophobic patch that is thought to help thioredoxin-based enzymes recognize and bind to unfolded proteins. It is possible that the hydrophobic patch helps to organize the hydrophobic core of I27 and accelerates folding, as is the case in other typical chaperone proteins (Saio et al., 2014). PDI simply acts as a placeholder so that when the protein collapses to the molten globule state, at 5.0 pN for example, the free thiol of titin can attack the mixed disulfide with PDI. With the reformed internal disulfide, the natively folded state is then quickly achieved. These studies demonstrate that even a substrate such as titin, which is under a constitutive load of no less than 4 pN in a resting muscle, can be successfully re-oxidized by a typical mammalian oxidoreductase enzyme.

These magnetic tweezers experiments further allow for the testing of key hypotheses related to titin protein folding during muscle contraction through direct comparison with equivalent single molecule assays performed on myosin II motors. Attempts to reconcile full muscle and muscle fiber mechanics experiment data with the trajectories observed from single molecule optical tweezers experiments on muscle myosin motors demonstrate the complexity of scaling mechanical systems over many orders of magnitude. Early single molecule optical tweezers experiments show that single myosin II power strokes generate steps of 12 nm and forces of 3.4 pN on average (Finer et al., 1994). Improvements such as using small ensembles of myosins to enable continuous sliding against a pulling force reported velocities of 500 nm/s at forces of 3 pN per motor (Debold et al., 2005), and the most recent experiments using entire myosin-actin cofilaments extracted from frog muscle observed unloaded velocities of 1,000 nm/s at 1 mM ATP and 20 pN (half the isometric force), providing a power output of 20,000 zW (Kaya et al., 2017).

Does scaling of the single molecule titin folding data to the level of the sarcomere explain some portion of the power output from muscle? The power measured from our eight Ig domain construct reaches a peak output of 6,000 zW at 4.0 pN, the force we have estimated titin to experience during a muscle contraction (Rivas-Pardo et al., 2016). Additionally, the power output from a real titin isoform, containing up to 100 tandem Ig domains in the I-band (Bang et al., 2001), would scale (according to equation (2)) to ~15,000 zW even if only 20% of the Ig domains were recruited to the unfolded state for the contraction. The premise advanced by Bianco and colleagues in their brief note therefore underestimates the contribution of titin to the output power of a muscle contraction by several orders of magnitude (Bianco et al., 2016). Scaling the 20,000 zW power output from single myosin-actin rod co-filaments of Kaya et al. to consider simultaneous interaction of the thick filament with six actin fibers (120,000 zW) plus the contribution from six parallel titin filaments (90,000 zW) provides a total power output per half sarcomere of 210,000 zW in accord with the estimate from Bianco, but demonstrating that a significant portion of the power (nearly half) if generated by titin folding.

This comparison between single molecule data on motors and folding Ig domains is central to the debate over their relative contributions to energy released during a muscle contraction. The magnetic tweezers data presented here demonstrates that a truncated chain of only eight disulfide-containing Ig domains are capable of folding at speeds of up to 1,900 nm/s, and power output of > 6,000 zW, which should renew the debate over how to incorporate titin dynamics into models of muscle contraction. Harnessing the chemical potential energy of ATP is clearly only one means by which animals power their motion; a hybrid mechanical system composed of titin and myosin in parallel can be tapped to deliver explosive levels of power through the synergy of protein folding and mechanochemical cycling of the myosin. It is therefore inescapable that the unique material properties of tandem protein domains, which we have demonstrated here, can used to power motion in heretofore undiscovered molecular and cellular systems, one of which happens to be muscle.

## Materials and Methods

### Instrumentation and protein purification

All single molecule magnetic tweezers experiments were performed as described previously, utilizing HaloTag chemistry for attachment to the glass and biotin-streptavidin for attachment to superparamagnetic beads(Popa et al., 2016; Rivas-Pardo et al., 2016). The octameric I27 DNA constructs were assembled as described previously (Wiita et al., 2006). I27^OXD^ was modified to contain an N-terminal SpyCatcher (Zakeri et al., 2012) and a C-terminal AviTag (Avidity). The protein was purified by Ni-NTA affinity and incubated overnight with 0.03% H_2_O_2_ to induce disulfide formation. Buffer was exchanged with PBS via dialysis for the biotinylation reaction, followed by gel filtration as described previously (Kosuri et al., 2012). The octamer protein was finally reacted with a HaloTag-SpyTag construct, purified via Ni-NTA affinity and gel filtration.

### Single molecule recordings

A solution containing ~10 nM I27^OXD^ was added to the flow cell for 20 minutes and was washed out with an experimental buffer consisting of PBS buffer pH 7.4 containing 10 mM ascorbate to prevent nonspecific oxidation of cysteine thiols and other side chains (Valle-Orero et al., 2017). Streptavidin beads blocked were blocked the previous night with 1% BSA in PBS, which was exchanged to the experimental PBS/ascorbate buffer immediately before the experiment. 50 uL of the streptavidin beads were flowed through the chamber and allowed to settle on the bottom coverslip, after which the permanent magnets were brought down to a resting position that applies 4 pN to the beads. Fresh experimental buffer was frequently exchanged into the flow cell, every ~30 minutes, due to evaporative losses from the open ends of the flow cell.

**Additional Materials and Methods can be found in the Supplementary Information.**

## Acknowledgements

This work was funded by the generous support of the National Institutes of Health grant R01HL61228 (to JMF) and fellowship F30HL129662 (to ECE).

